# Ultra-Fast Homomorphic Encryption Models enable Secure Outsourcing of Genotype Imputation

**DOI:** 10.1101/2020.07.02.183459

**Authors:** Miran Kim, Arif Harmanci, Jean-Philippe Bossuat, Sergiu Carpov, Jung Hee Cheon, Ilaria Chillotti, Wonhee Cho, David Froelicher, Nicolas Gama, Mariya Georgieva, Seungwan Hong, Jean-Pierre Hubaux, Duhyeong Kim, Kristin Lauter, Yiping Ma, Lucila Ohno-Machado, Heidi Sofia, Yongha Son, Yongsoo Song, Juan Troncoso-Pastoriza, Xiaoqian Jiang

## Abstract

Genotype imputation is a fundamental step in genomic data analysis such as GWAS, where missing variant genotypes are predicted using the existing genotypes of nearby ‘tag’ variants. Imputation greatly decreases the genotyping cost and provides high-quality estimates of common variant genotypes. As population panels increase, e.g., the TOPMED Project, genotype imputation is becoming more accurate, but it requires high computational power. Although researchers can outsource genotype imputation, privacy concerns may prohibit genetic data sharing with an untrusted imputation service. To address this problem, we developed the first fully secure genotype imputation by utilizing ultra-fast homomorphic encryption (HE) techniques that can evaluate millions of imputation models in seconds. In HE-based methods, the genotype data is end-to-end encrypted, i.e., encrypted in transit, at rest, and, most importantly, in analysis, and can be decrypted only by the data owner. We compared secure imputation with three other state-of-the-art non-secure methods under different settings. We found that HE-based methods provide full genetic data security with comparable or slightly lower accuracy. In addition, HE-based methods have time and memory requirements that are comparable and even lower than the non-secure methods. We provide five different implementations and workflows that make use of three cutting-edge HE schemes (BFV, CKKS, TFHE) developed by the top contestants of the iDASH19 Genome Privacy Challenge. Our results provide strong evidence that HE-based methods can practically perform resource-intensive computations for high throughput genetic data analysis. In addition, the publicly available codebases provide a reference for the development of secure genomic data analysis methods.

## Introduction

Whole-genome sequencing (WGS)^1,2^ has become the standard technique in clinical settings for tailoring personalized treatments^3^ and in research settings for building reference genetic databases^4–6^. Technological advances in the last decade enabled a massive increase in the throughput of WGS methods^7^, which provided the opportunity for population-scale sequencing^8^, where a large sample from a population is sequenced, for studying ancestry, complex phenotypes^9,10^, as well as rare^11–13^ and chronic diseases^14^. While the price of sequencing has been decreasing, the sample sizes are increasing to accommodate the power necessary for new studies. It is anticipated that tens of millions of individuals will have access to their personal genomes in the next few years. The massive increase in the number of sequenced genomes makes it very challenging to share and analyze these datasets. There is a very strong need for new methods and frameworks that will enable decreasing the cost and facilitate analysis and management of genome sequencing.

One of the main techniques used for decreasing the cost of large-scale genotyping is in-silico genotype imputation, i.e., measuring genotypes at a subsample of variants, e.g., using a genotyping array and then utilizing the correlations among the genotypes of nearby variants (the variants that are close to each other in genomic coordinates) and imputing the missing genotypes using the sparsely genotyped variants^15–17^. Imputation methods aim at capturing the linkage disequilibrium patterns on the genome. These patterns emerge because the genomic recombination occurs at hotspots rather than at uniformly random positions along the genome. The genotyping arrays are designed around the idea of selecting a small set of ‘tag’ variants that, as small as 1% of all variants, optimize the trade-off between the cost and the imputation accuracy^18,19^. Imputation methods learn the correlations among variant genotypes by using population-scale sequencing projects^20^. In addition to filling missing genotypes, the imputation process has many other advantages. Combining low-cost array platforms with computational genotype imputation methods decreases genotyping costs and increases the power of genome-wide association studies (GWAS) by increasing the sample sizes^21^. Accurate imputation can also greatly help with the fine mapping of causal variants^22^ and is vital for meta-analysis of the GWAS^23^. Genotype imputation is now a standard and integral step in performing GWAS. Although imputation methods can predict only the variant genotypes that exist in the panels, the panels’ sample sizes are increasing rapidly, e.g., in projects such as TOPMed^24,25^ will provide training data for imputation methods to predict rarer variant genotypes, and this can increase the sensitivity of GWAS.

Although imputation and sparse genotyping methods enable a vast decrease in genotyping costs, they are computationally very intensive and require management of large genotype panels and interpretation of the results^26^. The imputation tasks can be outsourced to third parties, such as the Michigan Imputation Server, where users upload the genotypes (as a VCF file) to a server that performs imputation internally using a large computing system. The imputed genotypes are then sent back to the user. There are, however, major privacy^27^ and data security^28^ concerns over using these services, since the genotype data is analyzed in plaintext format where any adversary who has access to the third party’s computer can view, copy, or even modify the genotype data. As genotype imputation is one of the central initial steps in many genomic analysis pipelines, it is essential that the imputation be performed securely to ensure that these pipelines can be computed securely as a whole. For instance, although several secure methods for GWAS have been developed^29^, as long as genotype imputation (a vital step in GWAS analyses) is not performed securely, it is almost nonsensical to analyze the security of GWAS analyses.

Genomic data security and privacy has received much attention in the recent years. Most notably, the increasing prevalence of genomic data, e.g., direct-to-consumer testing and recreational genealogy, make it harder to share genomic data due to privacy concerns. Genotype data is very accurate in identifying the owner because of its high dimensionality, and leakage can cause concerns about discrimination and stigmatization^30^. Also, the recent cases of forensic usage of genotype data are making it very complicated to share data for research purposes. The identification risks extend to family members of the owner since a large portion of genetic data is shared with relatives. Many attacks have been proposed on genomic data sharing models, where the correlative structure of the variant genotypes provides enough power to adversaries to make phenotype inference and individual identification^31^ possible. Therefore, it is of the utmost importance to ensure that genotype data is shared securely.

To address these growing genomic data security concerns, we developed and implemented several approaches for secure genotype imputation. Our methods make use of the homomorphic encryption (HE) formalism^32^ that provides mathematically provable, and potentially the strongest security guarantees for protecting genotype data while imputation is performed in an untrusted semi-honest environment. To include a comprehensive set of approaches, we focus on three state-of-the-art HE cryptosystems, namely Brakerski/Fan-Vercauteren (BFV)^33,34^, Cheon-Kim-Kim-Song (CKKS)^35^, and Fully Homomorphic Encryption over the Torus (TFHE)^36,37^. In our HE-based framework, genotype data is encrypted by the data owner before outsourcing the data. After this point, data remains always encrypted, i.e., encrypted in-transit, in-use, and at-rest; it is never decrypted until the results are sent to the data owner. The strength of our HE-based framework stems from the fact that the genotype data remains encrypted even while the imputation is being performed. Hence, even if the imputation is outsourced to an untrusted third party, any semi-honest adversaries learn nothing from the encrypted data. This property makes the HE-based framework very powerful: For an untrusted third party who does not have access to the private key, the genotype data is indistinguishable from random noise (i.e., practically of no use) at any stage of the imputation process. Our HE-framework provides the strongest form of security for outsourcing genotype imputation compared to any other approaches under the same adversarial model.

HE-based frameworks have been deemed impractical since their inception. Therefore, in comparison to other cryptographically secure methods, such as multiparty computation^29^ and trusted execution environments^38^, HE-based frameworks have received little attention. Recent theoretical breakthroughs in the HE literature, and a strong community effort^39^ have since rendered HE-based systems practical. Many of these improvements, however, are only beginning to be reflected in practical implementations and applications of HE algorithms. In this study, we provide evidence for the practicality of the HE formalism by building secure and ready-to-deploy methods for genotype imputation. We perform detailed benchmarking of the time and memory requirements of HE-based imputation methods and demonstrate the feasibility of large-scale secure imputation. In addition, we compared HE-based imputation methods with the state-of-the-art plaintext, i.e., non-secure, imputation methods, and we found comparable performance (with a slight decrease) in the imputation accuracy with the benefit of total genomic data security.

We present HE-based imputation methods in the context of two main steps, as this enables a general modular approach. The first step is imputation model building, where imputation models are trained using the reference genotype panel with a set of tag variants (variant genotypes on an Illumina array platform) to impute the genotypes for a set of target variants, e.g., common variants in the 1000 Genomes Project^5^ samples. The second step is the secure imputation step where the encrypted tag variant genotypes are used to predict the target genotypes (which are encrypted) by using the imputation models trained in the first step. This step, i.e., imputation model evaluation using the encrypted tag variant genotypes, is where the HE-based methods are deployed. In principle, the model training step needs to be performed only once when the tag variants do not change, i.e., the same array platform is used for multiple studies. Although these steps seem independent, model evaluation is heavily dependent on the representation and encoding of the genotype data, and the model complexity affects the timing and memory requirements of the secure outsourced imputation methods. Our results suggest, however, that linear models (or any other model that can be approximated by linear models) can be almost seamlessly trained and evaluated securely with almost no penalty on the performance, where the model builders (1st step) and model evaluators (2nd step) can work independently. We provide a pipeline that implements both model training and evaluation steps so that it can be run on any selection of tag variants. We make the implementations publicly available so that they can be used as a reference by the computational genomics community.

## Results

We present the scenario and the setting for secure imputation and describe the secure imputation approaches we developed. Next, we present accuracy comparisons with the current state-of-the-art non-secure imputation methods and the time and memory requirements of the secure imputation methods. Finally, we present the comparison of time and memory requirements of our secure imputation pipeline with the non-secure methods.

### Genotype Imputation Scenario

Figure 1a illustrates the secure imputation scenario. A researcher genotypes a cohort of individuals by using genotyping arrays or other targeted methods, such as whole-exome sequencing, and calls the variants using a variant caller such as GATK^40^. After the genotyping, the genotypes are stored in plaintext, i.e., unencrypted and not secure for outsourcing. Each variant genotype is represented by one of the three values {0, 1, 2}, where 0 indicates a homozygous reference genotype, 1 indicates a heterozygous genotype, and 2 indicates a homozygous alternate genotype. To secure the genotype data, the researcher generates two keys, a public key for encrypting the genotype data and a private key for decrypting the imputed data. The public key is used to encrypt the genotype data into ciphertext, i.e., random-looking data that contains the genotype data in a secure form. It is mathematically provable (i.e., equivalent to the hardness of solving the ring learning with errors, or RLWE, problem^41^) that the encrypted genotypes cannot be decrypted into plaintext genotype data by a third party without the private key, which is in the possession of only the researcher. Even if an unauthorized third party copies the encrypted data without authorization (e.g., hacking, stolen hard drives), they cannot gain any information from the data as it is essentially random noise without the private key. The security (and privacy) of the genotype data is therefore guaranteed, as long as the private key is not compromised. The security guarantee of the imputation methods is based on the fact that genotype data is encrypted in transit, during analysis, and at rest. The only plaintext data that is transmitted to the untrusted entity is the locations of the variants, i.e., the chromosomes and positions of the variants. This, however, does not lead to any security or privacy risks as the variant loci, for example on the genotyping arrays, are publicly known and do not provide any private information to an untrusted third party.

**Figure 1.**
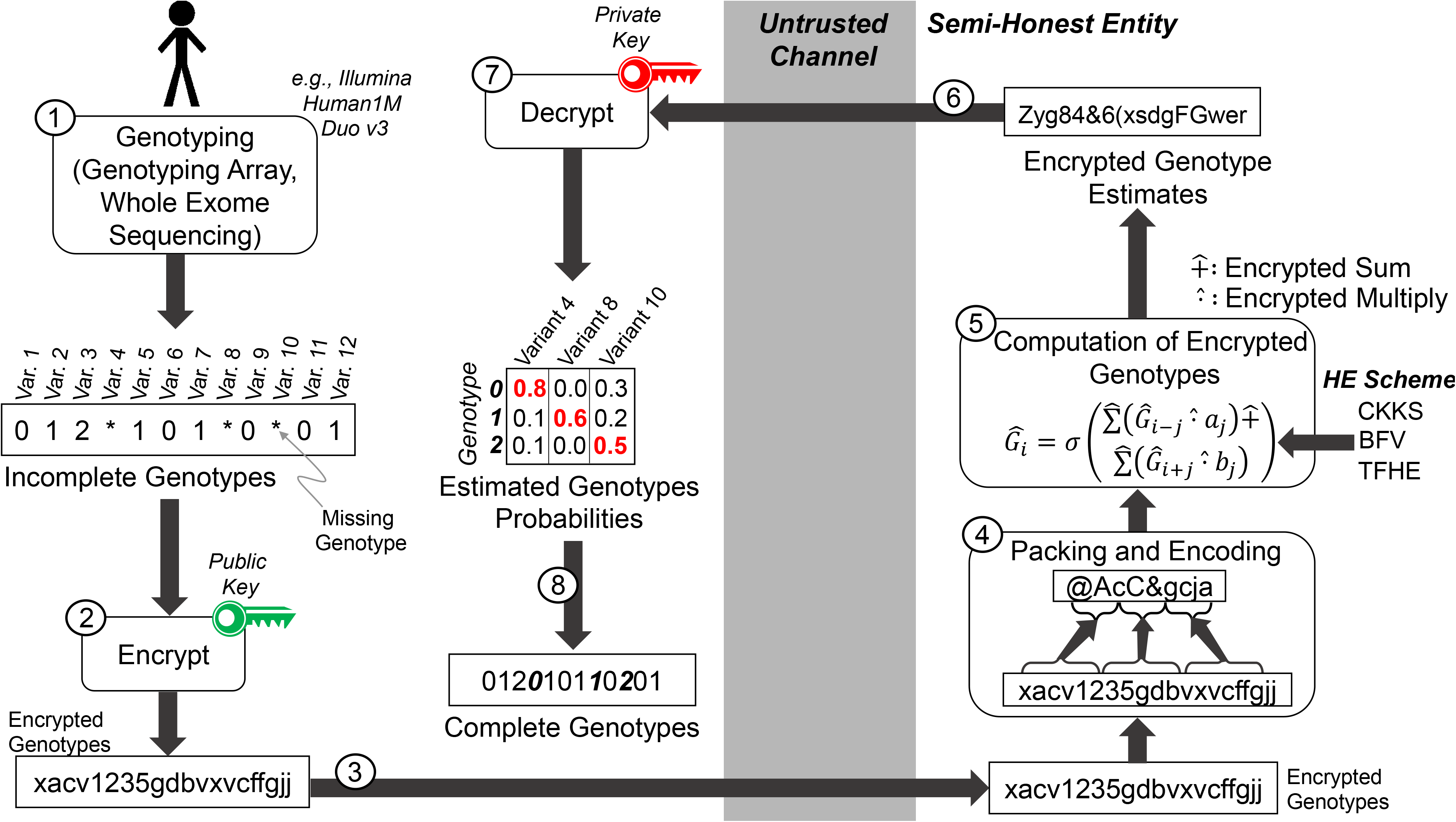
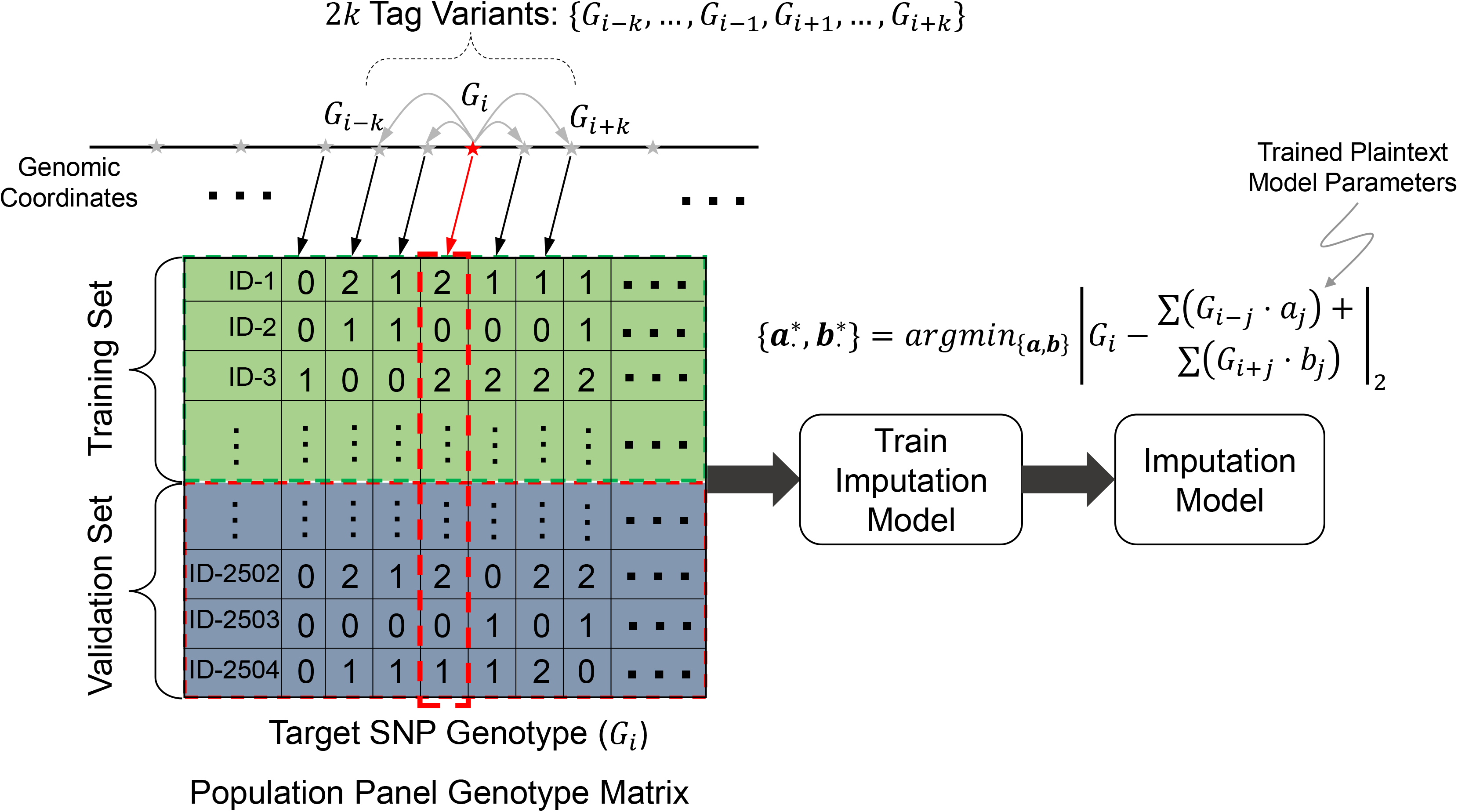
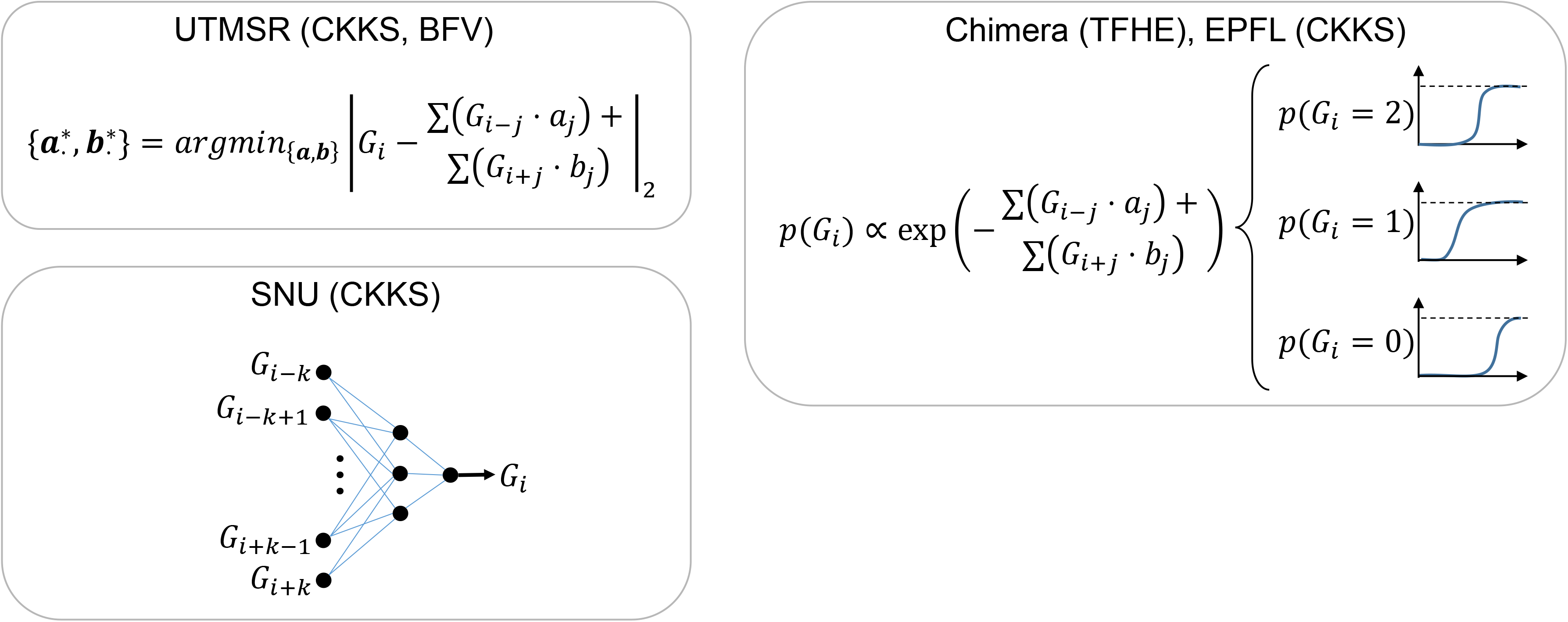
Illustration of secure genotype imputation. a) Illustration of the genotype imputation scenario. The incomplete genotypes are measured by genotyping arrays (e.g., Illumina Duo 1M Version 3) or whole-exome sequencing. 9 of the 12 variants are genotyped and 3 variants (variants 4, 8, and 10) have missing genotypes. The missing genotypes are represented by stars. The researcher encrypts the incomplete genotypes using the public key (Step 2). Encryption generates random-looking string from the genotypes. The encrypted genotypes are sent to the semi-honest entity, i.e., the imputation server (Step 3). The encrypted genotypes are first encoded and packed (Step 4) then they are input to the model evaluation (Step 5), which computes the estimates for the missing variant genotype probabilities. The estimated genotype probabilities are in the ciphertext domain and the semi-honest entity does not observe the genotype probabilities in plaintext. The encrypted genotype probabilities are then sent to the researcher (Step 6). The researcher decrypts the genotype probabilities (Step 7) and predicts the genotypes with highest probabilities and builds the final set of complete genotypes (Step 8). The imputed genotypes are indicated by bold and italicized font. b) Building of the plaintext model for genotype imputation. The semi-honest entity uses a publicly available population panel to build genotype estimation models for each variant. The panel is divided into training and validation samples set before the model is built. The training is performed using training samples and the accuracy is estimated using the validation set. The models are stored in plaintext domain. The model in the current study is a linear model where each variant genotype is modeled using genotypes of variants within a *k* variant vicinity of the variant, as illustrated on the top. c) The plaintext models implemented under the secure frameworks. UTMSR, Chimera, and EPFL teams use linear models. SNU team developed a neural network model with one hidden layer.

The encrypted genotypes are sent through a channel to the imputation service. The channel does not have to be secure against an eavesdropper because the genotype data is encrypted by the researcher. However, secure channels should regardless be authenticated to to prevent malicious man-in-the-middle attacks^42^. The encrypted genotypes are received by the imputation service, an honest-but-curious entity, i.e., they will receive the data legitimately and extract all the private information they can from the data. A privacy breach is, however, impossible as the data is always encrypted when it is in the possession of the imputation service. Hence, the only reasonable action for the secure imputation server is to perform the genotype imputation and to return the data to the researcher. It is possible that the imputation server acts maliciously and intentionally returns bad-quality data to the researcher, but there is no economically or academically reasonable motivation for such a malicious service, as it would be very easy to detect this behavior on the researcher’s side and to advertise the malicious or low quality of the service to other researchers. Therefore, we assume that the secure server is semi-honest, and it performs the imputation task as accurately as possible.

After receipt of the encrypted genotypes by the server, the first step is re-coding of the encrypted data into a packed format that is optimized for the secure imputation process. This step is performed to decrease time requirements and to optimize the memory usage of the imputation process. The data is coded to enable analysis of multiple genotypes in one cycle of the imputation process^43^. The next step is the secure evaluation of the imputation models, which entails securely computing the genotype probability for each variant by using the encrypted genotypes. The variants received from the researcher are treated as the tag variants whose genotypes are used as features in the imputation model to predict the “target” variants, i.e., the missing variants (Fig 1b). For each target variant, the corresponding imputation model uses the genotypes of the nearby tag variants to predict the target variant genotype in terms of genotype probabilities. In other words, we use a number of nearby tag variants to build an imputation model for the respective target variant such that the tag variants that are nearby (in genomic coordinates) are treated as the features for assigning genotype scores for the target variant. After the imputation is performed, the encrypted genotype probabilities are sent to the researcher. The researcher decrypts the genotype probabilities by using the private key. The final genotypes can be assigned using the maximum probability genotype estimate, i.e., by selecting the genotype with the highest probability for each variant.

As we describe in detail below, we developed a number of secure imputation methods, and we have developed, trained, and implemented numerous approaches for imputation. These approaches were selected because they performed very accurately in the iDASH19 Genomic Data Security Competition (Supplementary Information, Supplementary Table 1). In this study, we focus primarily on linear imputation models that can be trained very fast, using optimized libraries^44^. One of the key points of secure imputation is that the parameters of the imputation models can be stored in plaintext as they depend only on the population panels that are publicly available. Thus, the imputation server can train the imputation models in plaintext, i.e., training does not require secure computation. As the generalized linear models can be trained very rapidly and online, this enables a near real-time training potential in the case that the model parameters need to be trained when a new set of tag variants are used in the imputation. To utilize the plaintext model in the course of imputation, the additions and multiplications of the imputation model are converted to the ultra-fast and secure operations, where one of the operands (the genotype) is encrypted and the model parameters are plaintext (Fig. 1b, 1c). We focus on three different types of homomorphic encryption schemes, namely the BFV^33,34^, CKKS^35^, and TFHE^36^ schemes, and we provide implementations of the imputation models that use these schemes. We provide, in total, five approaches implemented by four different teams. For simplicity of description, we named these teams Chimera, EPFL, SNU, and UTMSR (See Methods). Among these, CKKS is used in three different approaches (UTMSR-CKKS, SNU-CKKS, EPFL-CKKS), and BFV and TFHE are each utilized by one separate approach (UTMSR-BFV, Chimera-TFHE, respectively). The teams independently developed plaintext imputation models, using different approaches, where Chimera team trained a logistic regression model and EPFL team trained a multinomial logistic regression model, both by using the nearby tag variants (Supplementary Figure 3, Table 7, and Tables 3,4); SNU used a 1-hidden layer neural network (Fig. 1c, Supplementary Figures 1, 2, Table 5); and UTMSR trained a linear regression model (Fig. 1c, Supplementary Figure 4). All the teams treat the genotypes as continuous predictions, except for Chimera and SNU teams who utilized a one-hot encoding of the genotypes (see Methods). The nearby variant selection is important as it determines how each target variant is imputed. In general, we found that the models that use 30-40 tag variants provide optimal results (for the current array platform) in terms of imputation accuracy (Supplementary Tables 2, 5, 6, 8). As previous studies have shown, we believe that tag variant selection can provide an increase in imputation accuracy^44^. Finally, we observed a general trend of linear scaling with the number of target variants (As shown in Supplementary Figure 5 and other supplementary tables). This provides evidence that there is no extra overhead (in addition to the linear increasing sample size) to scaling to genome-wide and population-wide computations.

### Accuracy Comparisons with the Non-Secure Methods

We first analyzed the imputation accuracy of the secure methods with the plaintext (non-secure) counterparts that are the most popular state-of-the-art imputation methods. We compared secure imputation methods with IMPUTE2^45^, Minimac3^46^, and Beagle^47^ methods. These plaintext methods utilize hidden Markov models (HMM) for genotype imputation (see Methods). The population panels and the pre-computed estimates of the recombination frequencies are taken as input to the methods. Each method is set to provide a measure of genotype probabilities, in addition to the imputed genotype values.

To perform comparisons in a realistic setting, we used the variants on the Illumina Duo 1M version 3 array platform^48^. This is a popular array platform that covers more than 1.1 million variants and is used by population-scale genotyping studies such as HAPMAP^49^. We extracted the genotypes of the variants that are probed by this array platform and overlap with the variants identified by the 1000 Genomes Project population panel of 2,504 individuals. For simplicity of comparisons, we focused on chromosome 22. The variants that are probed by the array are treated as the tag variants that are used to estimate the target variant genotypes. The target variants are defined as the variants on chromosome 22 whose allele frequency is greater than 5%, as estimated by the 1000 Genomes Project^5^. We used the 16,184 tag variants and 80,882 common target variants. Then, we randomly divided the 2,504 individuals into a training genotype panel of 1,500 samples and a testing panel of 1,004 samples. The training panel is used as the input to the plaintext methods (i.e., IMPUTE2, Minimac3, Beagle) and also for building the plaintext imputation models of the secure methods. Each method is then used to impute the target variants using the tag variants. Figure 2a shows the comparison of genotype prediction accuracy computed over all the predictions made by the methods. The non-secure methods show the highest accuracy among all the methods. The secure methods exhibit very similar accuracy, whereas the closest method follows with only a 2-3% decrease in accuracy. To understand the differences between the methods, we also computed the accuracy of the non-reference genotype predictions (see Methods, Fig. 2b). The non-secure methods show slightly higher accuracy compared to the secure methods. These results indicate that the proposed secure methods provide perfect data privacy at the cost of a slight decrease in imputation accuracy.

**Figure 2.**
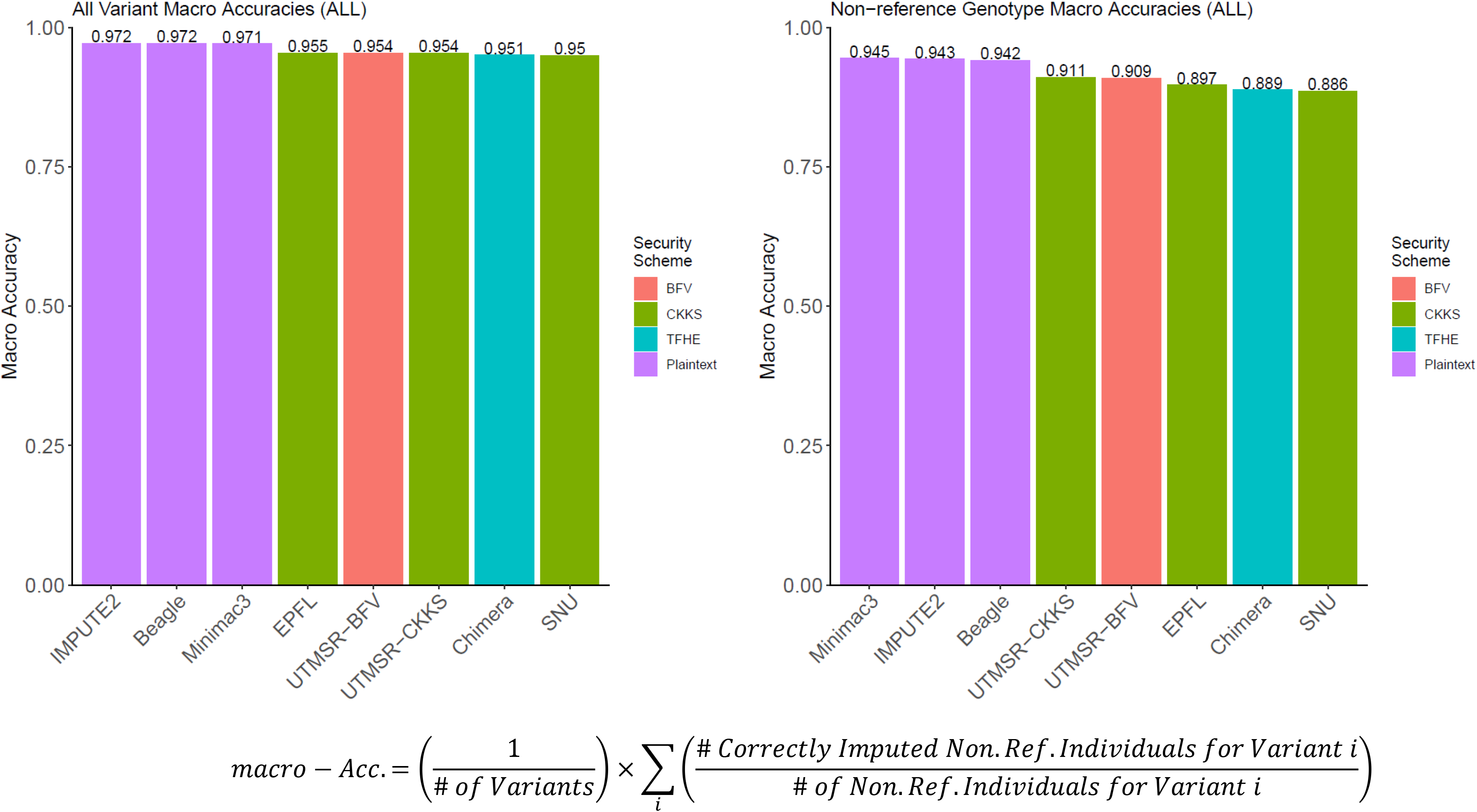
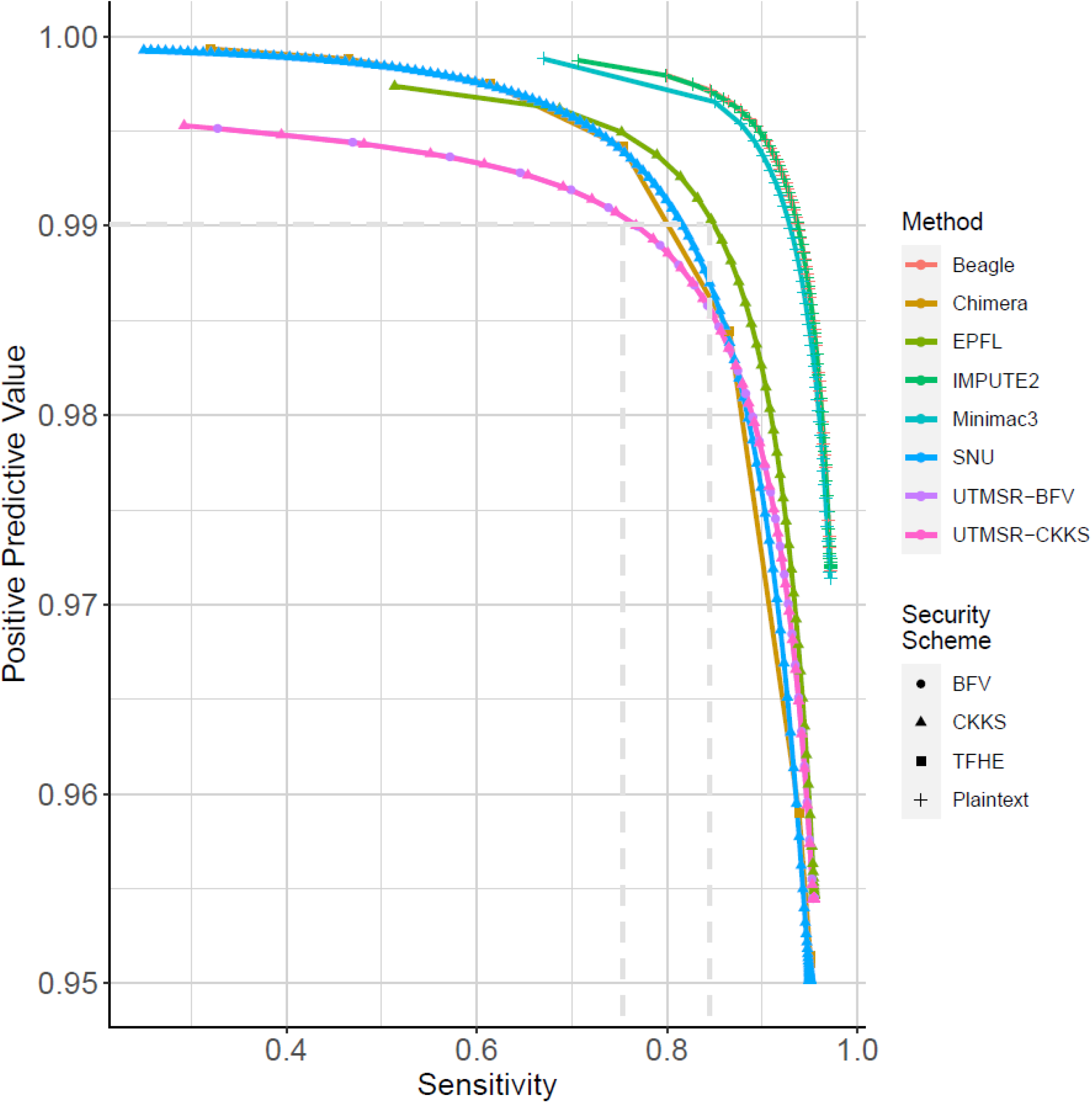
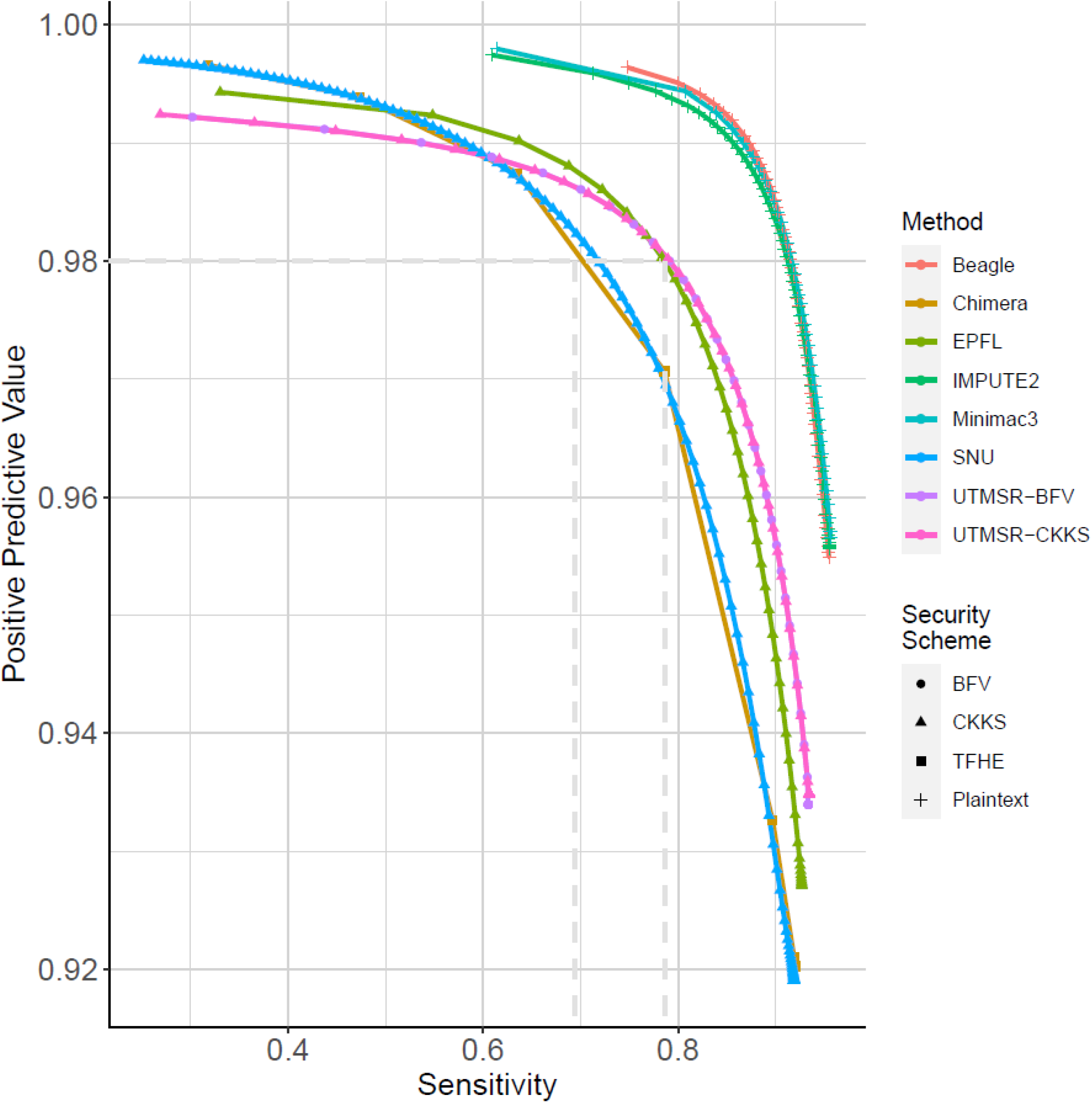
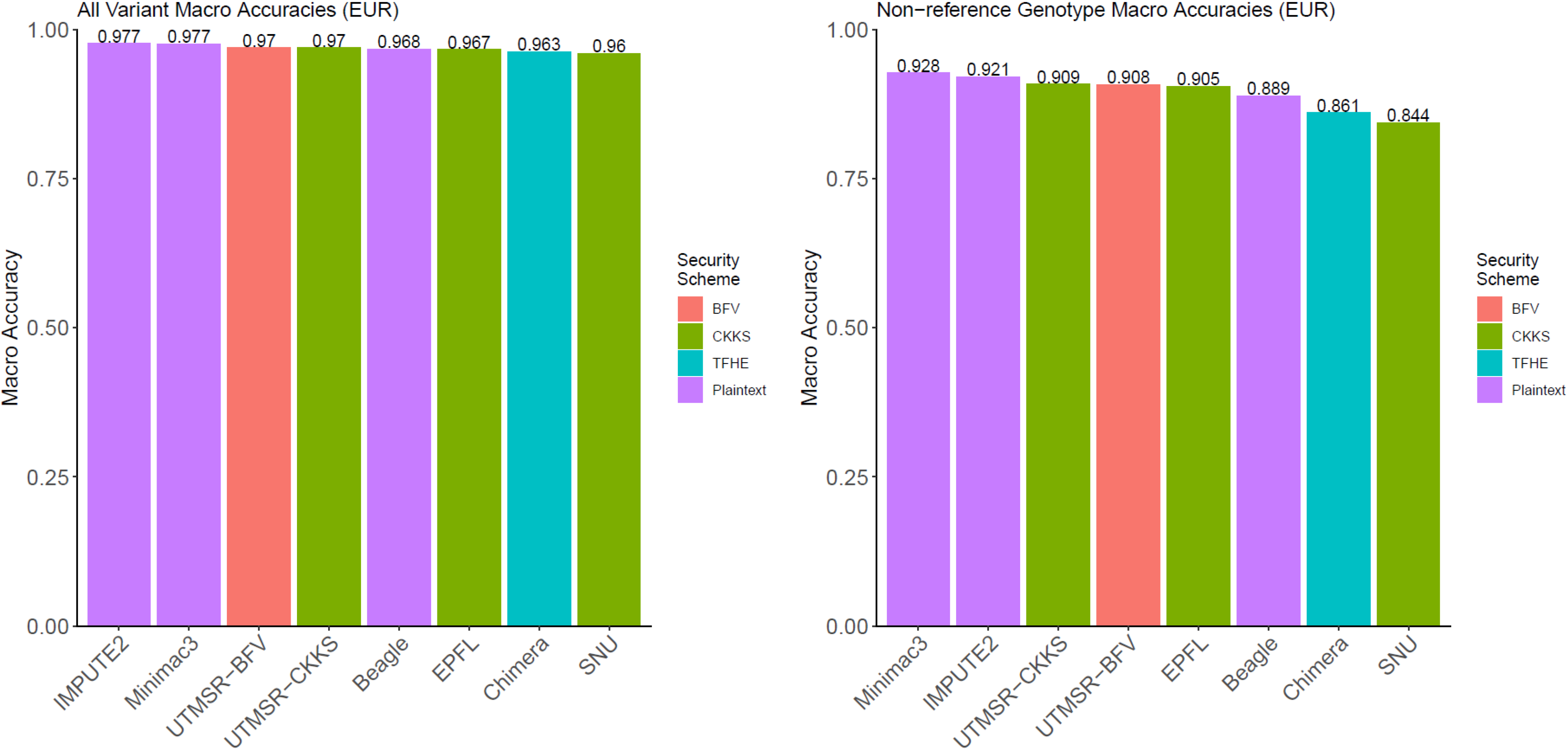
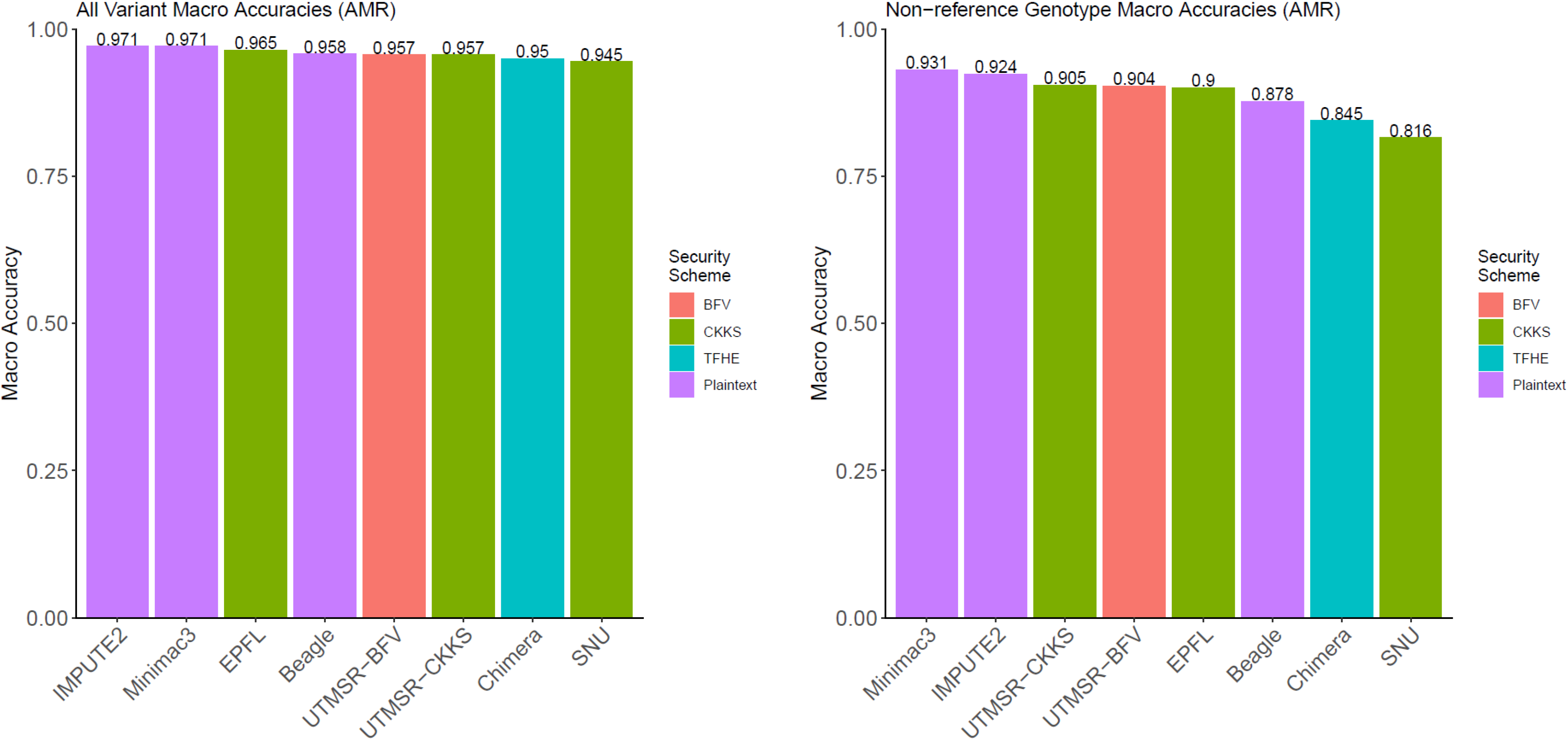
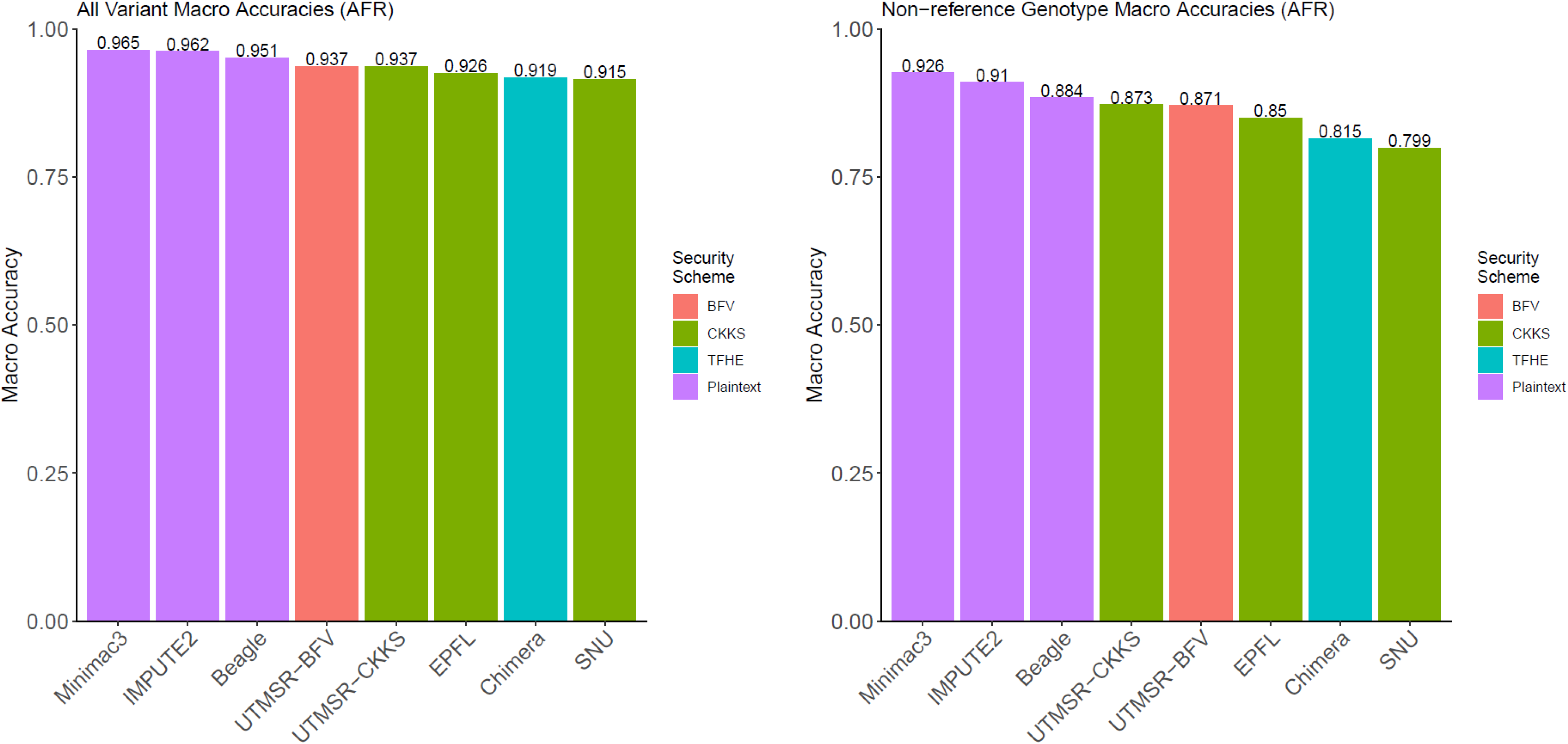
Accuracy comparisons with the non-secure methods. The accuracy of genotype imputation for all the genotypes (a) and the non-reference genotypes (b) are shown. The x-axis indicates the method name; the numeric accuracy value is included at the top of each bar for comparison. The sensitivity versus positive predictive value curves are plotted for more detailed accuracy comparisons for all genotypes (c) and the non-reference genotypes (d). Each curve corresponds to a method. The marker shapes indicate the type of security scheme that is implemented by each method. Plaintext indicates the non-secure methods. The population stratification of the accuracy is shown for EUR all genotypes (e) and non-ref genotypes (f), AMR all (g) and non-ref (h) genotypes, and AFR all (i), and non-ref genotypes (j).

We next assessed whether the genotype probabilities (or scores) computed from the secure methods provide meaningful measures for choosing reliably imputed genotypes. For this, we calculated the sensitivity and the positive predictive value (PPV) of the imputed genotypes whose scores exceed a cutoff (see Methods). To analyze how the cutoff selections affect the accuracy metrics, we shifted the cutoff (swept the cutoff over the range of genotype scores) so that the accuracy is computed for the most reliable genotypes (high cutoff) and for the most inclusive genotypes (low cutoff). We then plotted the sensitivity versus the PPV (Fig 2c). Compared to the secure methods, the non-secure methods generally show a higher sensitivity at the same PPV. However, secure methods can capture more than 80% of the known genotypes at 98% accuracy. The same results hold for the non-reference genotypes’ prediction accuracy (Fig 2d). These results indicate that secure genotype predictions can be filtered by setting cutoffs to improve accuracy.

We also evaluated the population-specific effects on the imputation accuracy. For this, we divided the testing panel into three populations, 210 European (EUR), 135 American (AMR), and 272 African (AFR) samples, as provided by the 1000 Genomes Project. The training panel yielded 389 AFR, 212 AMR, and 293 EUR samples. Figures 2e and 2f show genotype and non-ref genotype accuracy for EUR, respectively. We observed that the non-secure and secure methods are similar in terms of accuracy. We observed that the secure CKKS (UTMSR-CKKS) scheme with a linear prediction model outperformed Beagle in EUR population, with marginally higher accuracy. We observed similar results for AMR populations where the non-secure methods performed at the top and secure methods show very similar but slightly lower accuracy (Fig. 2g, 2h). For AFR populations, the non-reference genotype prediction accuracy is lower for all the methods (Fig. 2i, 2j). This is mainly rooted at the fact that the African populations show distinct properties that are not yet well characterized by the 1000 Genomes Panels. We expect that the larger panels can provide better imputation accuracy.

To further investigate the nature of the imputation errors, we analyzed the characteristics of imputation errors of each method by computing the confusion matrices (Supplementary Fig. 6). We found that the most frequent errors are made when the real genotype is heterozygous, and the imputed genotype is a homozygous reference genotype. The pattern holds predominantly in secure and non-secure methods, although the errors are slightly lower, as expected, for the non-secure methods. Overall, these results indicate that secure imputation models can provide genotype imputations comparable to non-secure counterparts.

### Timing and Memory Requirements of Secure Imputation Methods

One of the main unjustified critiques of homomorphic encryption methods is that they are impractical due to memory and time requirements. We, therefore, believe that the most important challenge is to make HE methods practical in terms of memory and time. To assess and demonstrate the practicality of the secure methods, we performed a detailed analysis of the time and memory requirements of secure imputation methods. We divided the imputation process into four steps (key generation, encryption, secure model evaluation, and decryption), and we measured the time and the overall memory requirements. Figure 3a shows the detailed time requirements for each step. In addition, we studied the scalability of secure methods. For this, we report the time requirements for 20,000 (20K), 40,000 (40K), and 80,000 (80K) target variants to present how the time requirements scale with the number of target variants. The secure methods spend up to 10 milliseconds for key generation. In the encryption step, all methods were well below 2 seconds. The most time-consuming step of evaluation took less than 10 seconds, even for the largest set of 80K variants. Decryption, the last step, took less than 2 seconds. Except for the key generation and encryption, all methods exhibited a linear scaling with the increasing number of target variants. Overall, the total time spent in secure model evaluation took less than 25 seconds (Fig. 3b). This could be ignored when compared to the total time requirements of the non-secure imputation. Assuming that time usage scales linearly with the number of target variants (Fig. 3a), 4 million variants can be 312 microseconds per variant per 1000 individuals ((25 sec × 1000 individuals)/(80, 000 variants × 1004 individuals)). It can evaluated in approximately 1,250 seconds, which is less than half an hour. In other terms, secure evaluation is approximately be decreased even further by scaling to a higher number of CPUs (i.e., cores on local machines or instances on cloud resources). In terms of memory usage, all methods required less than 15 gigabytes of main memory, and three of the five approaches required less than 5 gigabytes (Fig. 3c). These results highlight the fact that secure methods could be deployed on even the commodity computer systems.

**Figure 3.**
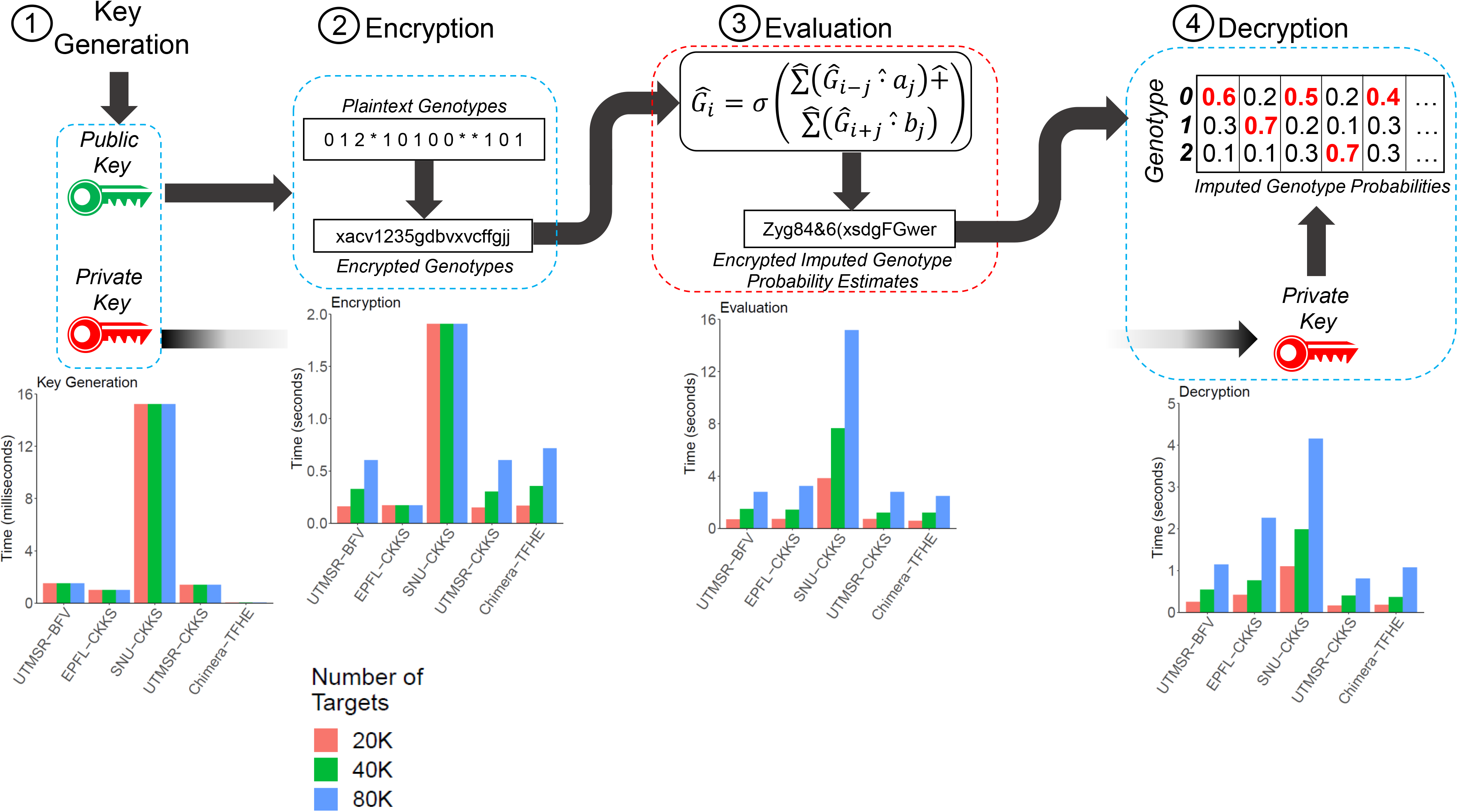
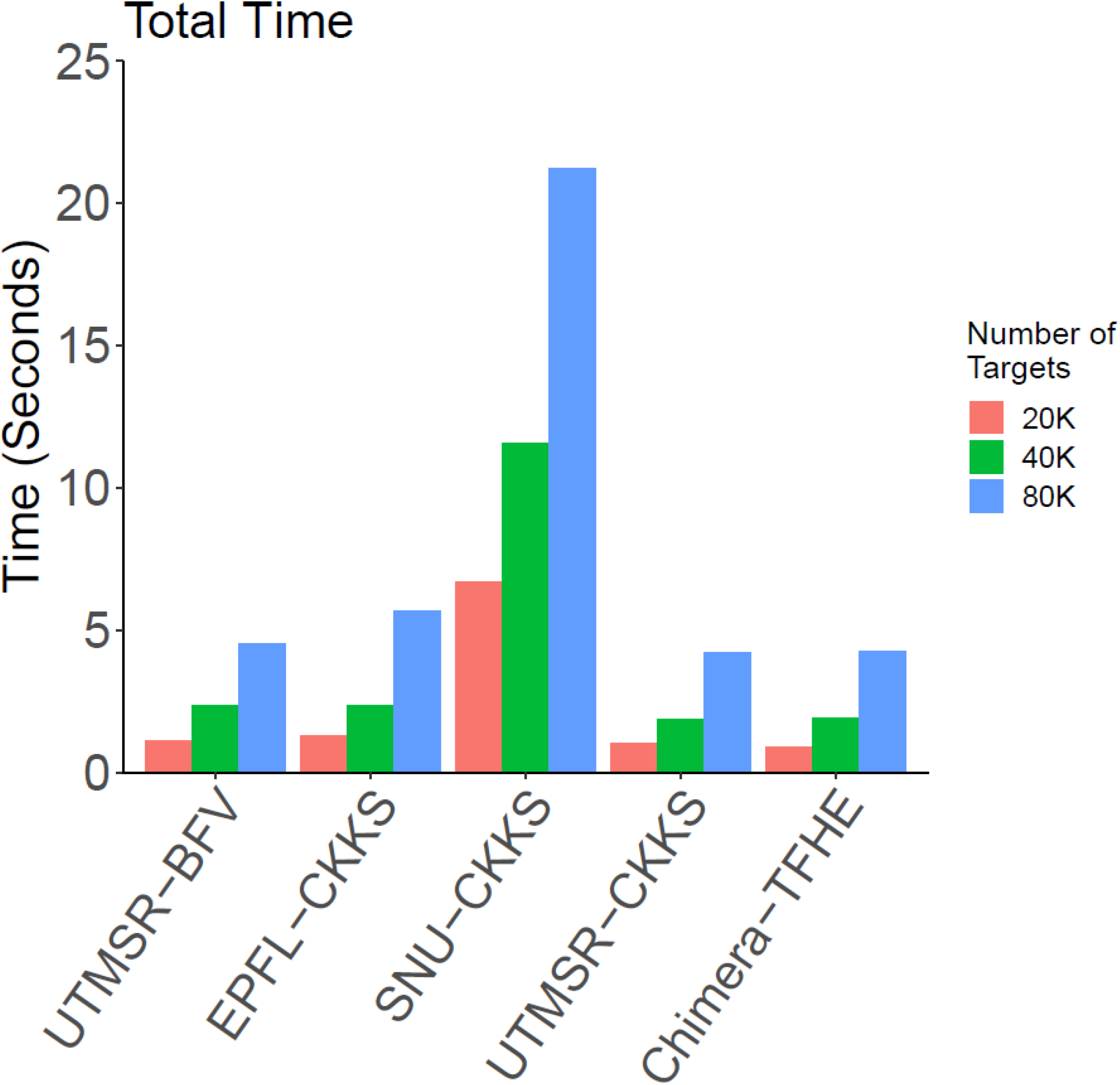
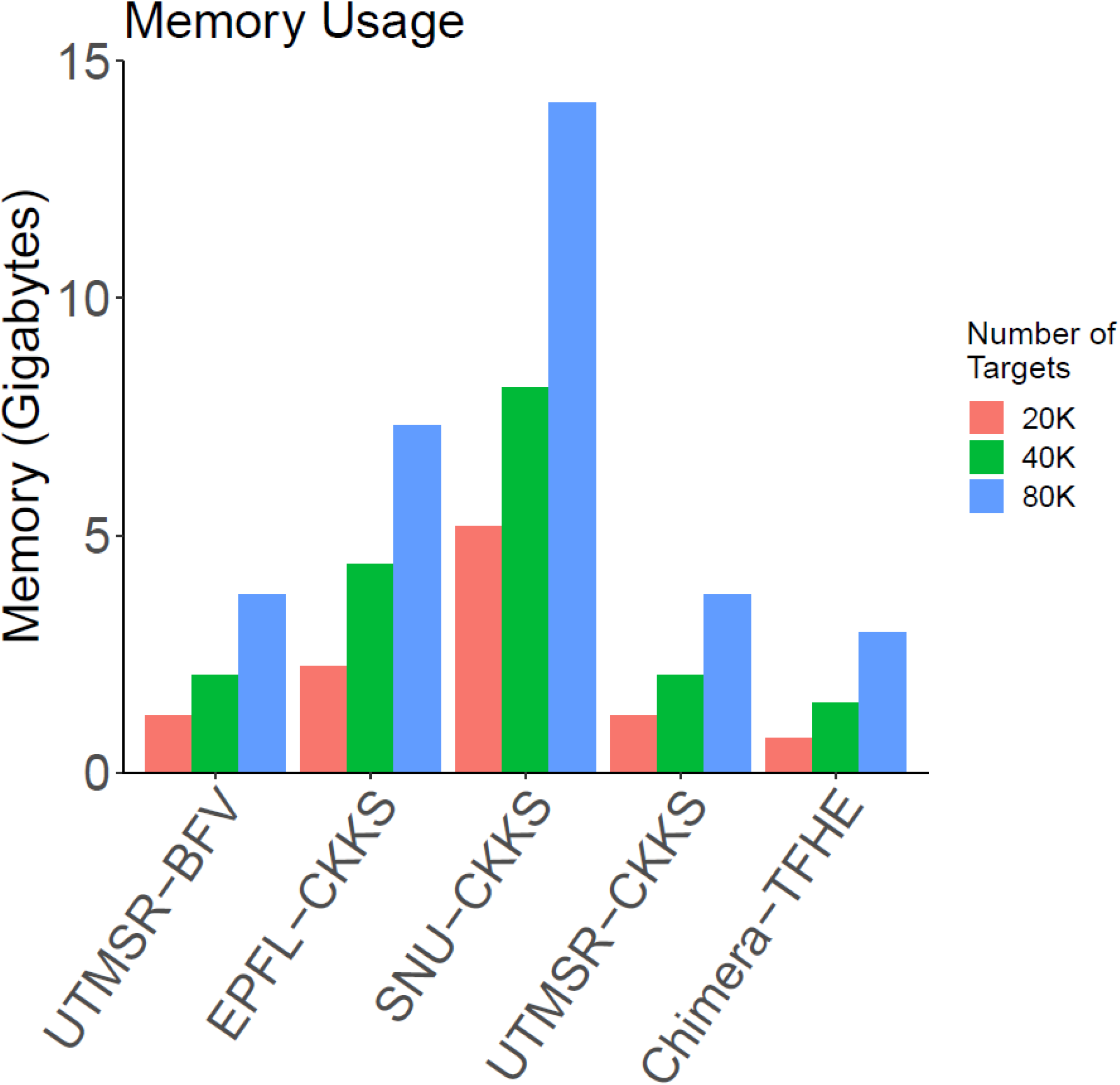
Detailed memory and time requirements of the secure methods. Each method is divided into 4 steps, (1) key generation, (2) Encryption, (3) Evaluation, (4) Decryption and each step is illustrated on the top of the figure in order. The cyan colored rectangles indicate the steps that are accomplished by the researcher and the red colored rectangle indicates the step 3 that is performed by the imputation service. The bar plots show the time requirements (a). Each step is performed using 20K, 40K, and 80K target variants to analyze the scaling of resource requirements. Bar colors indicate the input target variant sizes as specified in the legend. The aggregated time (b) and the maximum memory usage of the methods are also shown (c).

### Resource Usage Comparison between Secure and Non-Secure Imputation Methods

An important aspect of practicality is whether the methods are adaptable to different tag variants. This issue arises when a new array platform is used for genotyping the tag variants with a new set of tag variant loci. In this case, the current security framework requires that the plaintext models must be re-parametrized, and this may require a large amount of time. To evaluate this, we optimized the linear models for the UTMSR-CKKS approach and measured the total time (training + evaluation) and the memory for the 80K target variant set. This way, we believe that we perform a fair comparison of resource usage with other non-secure methods. We assumed that the training and secure evaluation would be run sequentially, and we measured the time requirement of the secure approach by summing the time for secure evaluation, assuming a maximum of 25 seconds from previous results, and the time for training. For memory, we computed the peak memory required for training and the peak memory required for secure evaluation. These time and memory requirements provided us with an estimate of resources used by the secure pipeline (Fig. 4a) that can be fairly compared to the non-secure methods.

**Figure 4.**
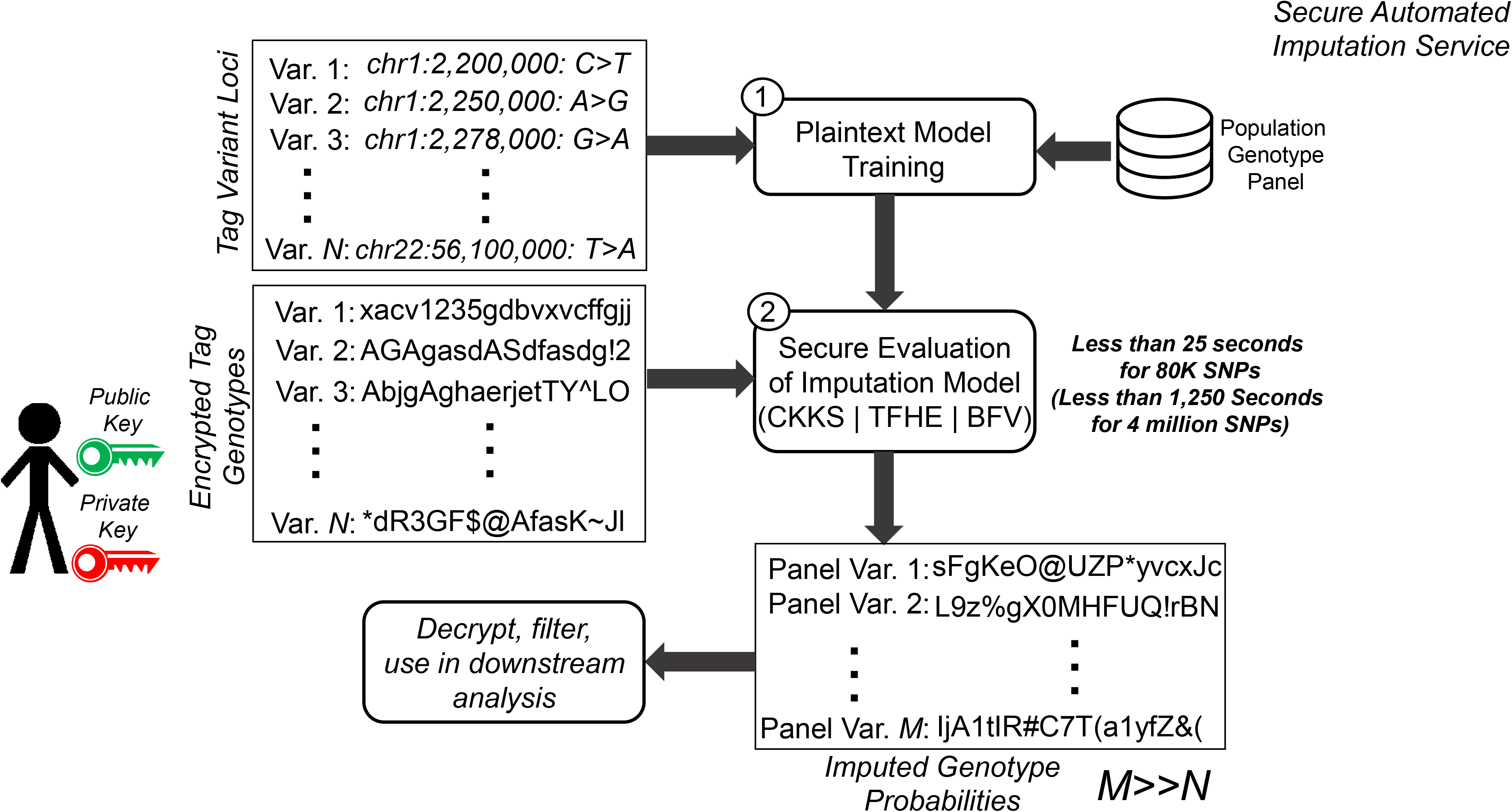
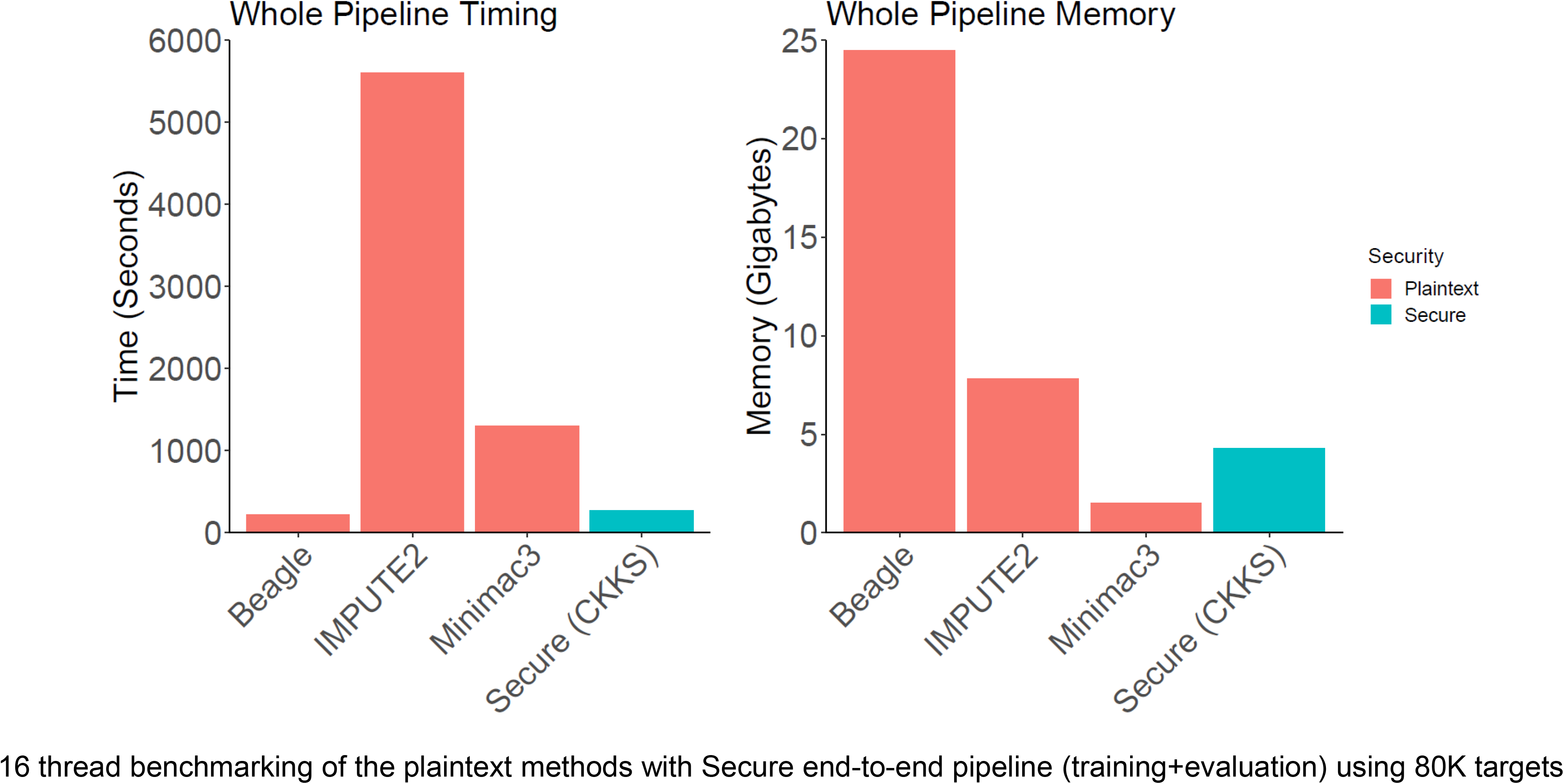
Illustration of a secure outsourced imputation service. The main steps of the secure service are the building of the plaintext model and evaluation of the model. The figure highlights the steps in the secure service where the researcher sends the variant loci to the service in the first step and the imputation service builds the plaintext imputation models using the publicly available population datasets. In the second step, the plaintext models are passed to the secure evaluation where one of the BFV, CKKS, or TFHE is used to securely estimate the imputed variant genotype probabilities. The final probabilities are sent back to the researcher. The random-looking strings represent the encrypted genotype data that cannot be decoded by anyone but the researcher. The time (b) and memory requirements (c) are illustrated in the bar plots. The color of the bars indicates the security scheme that is used by the methods. The y-axis shows the time (in seconds) and main memory (in gigabytes) used by each method to perform the imputation of the 80K variants where the secure outsourced method includes the plaintext model training and secure model evaluation steps.

We measured the time and memory requirements of all the methods by using a dedicated computer cluster to ensure resource requirements are measured accurately (see Methods). For IMPUTE2, there was no option for specifying multiple threads. Hence, we divided the sequenced portion of the chromosome 22 into 16 regions and imputed variants in each region in parallel using IMPUTE2, as instructed by the manual, i.e., we ran 16 IMPUTE2 instances in parallel to complete the computation. We then measured the total memory required by all 16 runs and used this as the memory requirement by IMPUTE2. We used the maximum time among all the 16 runs, as the time requirements by parallelized IMPUTE2. Beagle and Minimac3 were run with 16 threads as this option was available in the command line. Figures 4b and 4c show the time and memory requirements, respectively, of the three non-secure approaches and our secure method. The results show that the secure pipeline provides competitive timing (2nd fastest after Beagle) and memory requirements (2nd in terms of least usage after Minimac3). Our results also show that Minimac3 and our secure approach provided the best tradeoff between memory and timing, because Beagle and IMPUTE2 exhibit the highest time or highest memory requirements compared to other methods.

## Discussion

We presented fully secure genotype imputation methods that can practically scale to genome-wide imputation tasks by using ultra-fast homomorphic encryption techniques where the data is encrypted in transit, in analysis, and at rest. Unlike other approaches, such as the Michigan Imputation Server^50^ where the data is encrypted only at rest, our approach enables end-to-end encryption that is provably secure at every step of imputation. This is a unique aspect of the HE-based frameworks because, when appropriately performed, encryption is one of the few approaches that are recognized at the legislative level as a way of secure sharing of biomedical data, e.g. by HIPAA^51^ and partially by GDPR^52^. Our benchmarks with the state-of-the-art non-secure methods show that our pipelines can provide perfect genomic data security with very similar, or slightly lower, accuracy. Furthermore, our approaches have comparable and sometimes favorable time and memory requirements that exhibit practicality, even on commodity hardware. There are certain instances where the genomic data cannot be shared with a third party (e.g., data linkable to sensitive phenotypes, or consent is not provided). Our methods can offer a safe way to efficiently and practically outsource the compute-intensive and central task of genotype imputation.

Our study was enabled by several key developments in the fields of genomics and computer science. First, the recent theoretical breakthroughs in the homomorphic encryption techniques have enabled massive increases in the speed of secure algorithms. As much of the data science community still regards HE as a theoretical and not-so-practical framework, the reality is far from this image. We hope that our study can provide a reference for the development of privacy-aware and fully secure approaches that employ homomorphic encryption. Second, the amount of genomic data has increased several orders of magnitude in recent years. This provides enormous genotype databases where we can train the imputation models and test them in detail, before implementing them in secure evaluation frameworks. Another significant development is the recent formation of genomic privacy communities and alliances, i.e. Global Alliance for Genomic Health (GA4GH), where researchers build interdisciplinary approaches for developing privacy-aware methods. For example, our international study stemmed from the 2019 iDASH Genomic Privacy Challenge. We firmly believe that these communities will help bring together further interdisciplinary collaborations for the development of secure genomic analysis methods.

The presented imputation methods train an imputation model for each target variant. Our approaches handle millions of models, i.e., parameters. Unlike the HMM models that can adapt seamlessly to a new set of tag variants (i.e., a new array platform), our approaches need to be retrained when the tag variants are updated. We expect that the training can be performed a-priori for a new genotyping array and that they can be re-used in the imputation. The decoupling of the (1) plaintext training and (2) secure evaluation steps is very advantageous because plaintext training can be independently performed at the third party without the need to wait for the data to arrive. This way, the users would have to accrue only the secure evaluation time that is, as our results show, much smaller compared to the time requirements of the non-secure models, as small as 312 microseconds per variant per 1000 individuals. Nevertheless, even with the training, our results show that the secure imputation framework can train and evaluate in run times comparable to the plaintext (non-secure) methods. In the future, we expect many optimizations can be introduced to the models we presented. For example, we foresee that the linear model training can be replaced with more complex feature selection and training methods. Deep neural networks are potential candidates for imputation tasks as they can be trained for learning the complex haplotype patterns to provide better imputation accuracy^16^. With the introduction of the graphical processing units (GPUs) on the cloud, these models can be trained and evaluated securely and efficiently.

Finally, we believe that the multitude of models and the secure evaluation approaches that we presented here can help provide a much-needed reference point for the development and improvement of the imputation methods. Moreover, the developed models can be easily adapted to solve other privacy-sensitive problems by using secure linear and network model evaluations, such as the secure rare variant association tests^53^. Therefore, we believe that our codebases represent an essential resource for the computational genomics community.

## Online Methods

We present and describe the data sources, accuracy metrics, and non-secure imputation method parameters. The detailed methods are presented in the Supplementary Information.

### Variant and Genotype Datasets

All the tag and target variant loci, and the genotypes are collected from the public resources. We downloaded the Illumina Duo 1M version 3 variant loci from the array’s specification at the Illumina web site (https://support.illumina.com/downloads/human1m-duo_v3-0_product_files.html). The file was parsed to extract the variants on chromosome 22, which yielded 17,777 variants. We did not use the CNVs and indels while filtering the variants and we focused only on the single nucleotide polymorphims (SNPs). We then intersected these variants with the 1000 Genomes variants on chromosome 22 to identify the array variants that are detected by the 1000 Genomes Project. We identified 16,184 variants from this intersection. This variant set represents the tag variants that are used to perform the imputation. The phased genotypes on chromosome 22 for the 2,504 individuals in the 1000 Genomes Project are downloaded from the NCBI portal (ftp://ftp-trace.ncbi.nih.gov/1000genomes/ftp/release/20130502/ALL.chr22.phase3_shapeit2_mvncall_integrated_v5a.20130502.genotypes.vcf.gz). We filtered out the variants for which the allele frequency reported by the 1000 Genomes Project is less than 5%. After excluding the tag variants on the array platform, we identified 83,072 target variants that are to be used for imputation. As the developed secure methods use vicinity variants, the variants at the ends of the chromosome are not imputed. We believe this is acceptable because these variants are located very close to the centromere and at the very end of the chromosome. After filtering the non-imputed variants, we focused on the 80,882 variants that were used for consistent benchmarking of all the secure and non-secure methods.

### Accuracy Benchmark Metrics

We describe the genotype level and variant level accuracy. For each variant, we assign the genotype with the highest assigned genotype probability. The variant level accuracy is the average variant accuracy where each variant’s accuracy is estimated based on how well these imputed genotypes of the individuals match the known genotypes:

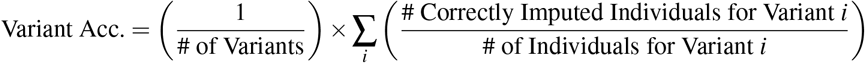

Variant level accuracy is also referred to as the macro-aggregated accuracy.

At the genotype level, we simply count the number of correctly computed genotypes and divide this with the total number of genotypes:

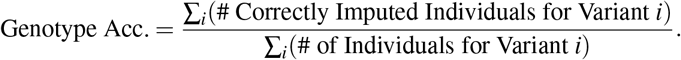

In the sensitivity vs positive predictive value (PPV) plots, the sensitivity and PPV are computed after filtering the imputed genotypes with respect to the imputation probability. We compute the sensitivity at the probability cutoff of *τ* is:

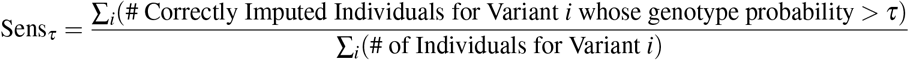

Positive predictive value measures the fraction of correctly imputed genotypes among the genotypes whose probability is above the cutoff threshold:

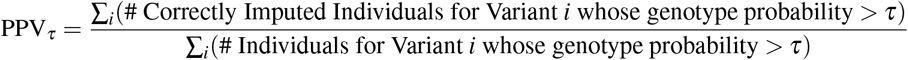

Next, we swept a large cutoff range for *τ* from −5 to 5 with steps 0.01. We finally plotted the sensitivity versus PPV to generate the precision-recall curves for each method.

#### Micro-AUC Accuracy Statistics

For parametrizing the accuracy and demonstrating how different parameters affect algorithm performance, we used micro-AUC as the accuracy metric. This was also the original accuracy metric for measuring the algorithm performance in iDASH19 competition. Micro-AUC treats the imputation problem as a three-level classification problem where each variant is “classified” into one of three classes, i.e., genotypes, {0, 1, 2}. Micro-AUC computes an AUC metric for each genotype then microaggregates the AUCs for all the genotypes. This enables assigning one score to a multi-class classification problem. We use the implementation in scikit-learn package to measure the micro-AUC scores for each method (https://scikit-learn.org/stable/modules/generated/sklearn.metrics.roc_auc_score.html).

#### Measurement of Time and Memory Requirements

For consistently measuring the time and memory usage among all the benchmarked methods, we used `“/usr/bin/time -f %e “\t” %M”` to report the wall time (in seconds) and peak memory usage (in kilobytes) of each method.

### Secure Methods

We briefly describe the secure methods.

#### UTMSR-BFV and UTMSR-CKKS

The UTMSR (UTHealth-Microsoft Research) team uses a linear model with the nearby tag variants as features for each target variant. The plaintext model training is performed using the GSL library. The collinear features are removed by performing the singular value decomposition (SVD) and removing features with singular values smaller than 0.01. The target variant genotype is modeled as a continuous variable that represents the “soft” estimate of the genotype (or the estimated dosage of the alternate allele), and can take any value from negative to positive infinity. The genotype probabilities are assigned by converting the soft genotype estimation to a score in the range [0,1]:

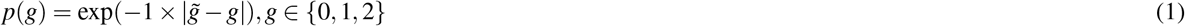

where *g* denotes one of the genotypes and 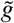 represents the decrypted value of the imputed genotype estimate. Suppose that each variant genotype is modeled using genotypes of variants within *k* variant vicinity of the variant. In plaintext domain, the imputed value can be written as follows:

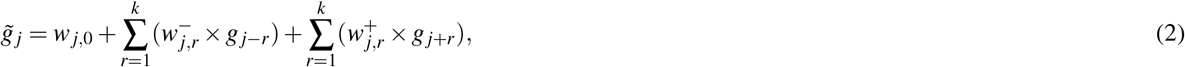

where *w*_*j*,0_ is the intercept of the linear model, and 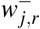 and 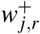 denote the linear model weights for the *j*^*th*^ target variant’s *r*^*th*^ upstream and downstream tag variants, respectively.

The secure outsourcing imputation protocols are implemented on two popular ring-based HE cryptosystems - BFV^33,34^ and CKKS^35^. These HE schemes share the same parameter setup and key-generation phase but have different algorithms for message encoding and homomorphic operations. In a nutshell, a ciphertext is generated by adding a random encryption of zero to an encoded plaintext, which makes the ring-based HE schemes secure under the RLWE assumption. More precisely, each tag variant is first encoded as a polynomial with its coefficients, and the encoded plaintext is encrypted into a ciphertext using the underlying HE scheme. The plaintext polynomial in the BFV scheme is separated from an error polynomial (inserted for security), whereas the plaintext polynomial in the CKKS scheme embraces the error. Then Eq. (2) is homomorphically evaluated on the encrypted genotype data by using the plain weight parameters. We exploit parallel computation on multiple individual data, and hence it enables us to obtain the predicted genotype estimates over different samples at a time. Our experimental results indicate that the linear model with 32 tag variants as features for each target variant shows the most balanced performance in terms of timing and imputation accuracy in the current testing dataset (see Supplementary Table 8 and Supplementary Figure 4). Our protocols achieve at least 128-bit security level from the HE standardization workshop paper^54^. We defer the complete details to the “UTHealth-Microsoft Research team solution” section in the Supplementary document.

#### Chimera-TFHE

The Chimera team used multi-class logistic regression (logreg) models trained over one-hot encoded tag features: each tag SNP variant is mapped to 3 Boolean variables. Chimera’s model training and architecture performed the best (with respect to accuracy and resource requirement) among six other solutions in the iDASH2019 Genotype Imputation Challenge.

We build three models per target SNP (one model per variant), i.e., target SNPs are also one-hot-encoded. These models give the probabilities for each target SNP variant. The maximal probability variant is the imputed target SNP value. A fixed number *d* of the nearest tag SNPs (in relation to the current target SNP) are used in model building. We train the models with different values of *d* in order to study the influence of neighborhood size: from 5 to 50 neighbors with an increment of 5. The most accurate model, in terms of micro-AUC score, is obtained for a neighborhood size *d* = 45. The fastest model with an acceptable accuracy (micro-AUC > 0.99) is obtained for *d* = 10. Although, the execution time of the fastest model is only ≈ 2 times faster compared to the most accurate model (refer to Table 2 in the Supplementary document).

During the homomorphic evaluation, only the linear part of the logreg model is executed, which means in particular that we do not homomorphically apply the sigmoid function on the output scores. We use the coefficient packing strategy and pack as many plaintext values as possible in a single ciphertext. The maximum number of values that can be packed in a RingLWE ciphertext equals the used ring dimension, which is *n* = 1024 in our solution. We chose to pack one or several columns of the input (tag SNPs) into a single ciphertext. Since the TFHE library RingLWE ciphertexts encrypt polynomials with Torus 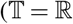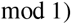 coefficients, we downscale the data to Torus values (multiples of 2^−14^) and upscale the model coefficients to integers.

In our solution, we use linear combinations with public integer coefficients. The evaluation is based on the security of LWE and only the encryption phase uses RingLWE security notions with no additional bootstrapping or key-switching keys. The security parameters have been tuned to support binary keys. Of course, as neither bootstrapping nor key-switching is used in our solution, the key distribution can be changed to any distribution (including the full domain distribution) without any time penalty. Our scheme achieves 130 bits of security, according to the LWE estimator^55^. More information about the our solution is described in the supplementary document (“Chimera-TFHE team solution”).

#### EPFL-CKKS

EPFL uses a multinomial logistic regression model with *d* − 1 neighboring coefficients and 1 intercept variable for each target variant, with three classes {0,1,2}. The plaintext model is trained using the `scikit-learn` python library. The input variants are represented as values {0,1,2}. There is no pre-processing applied to the training data. For a target position *j*, the predicted probabilities for each class label are given by:

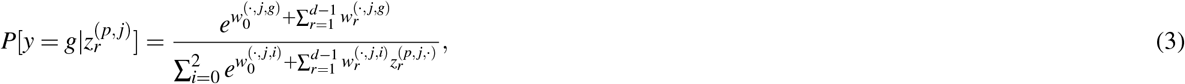

where 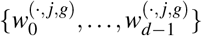 are the trained regression coefficients for label *g* ∈ {0, 1, 2} and position *j*, and 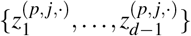 are the neighboring variants for patient *p* around target position *j*. The hard prediction for position *j* is given by 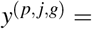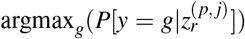. The variants 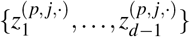 are sent encrypted and packed to the server, using the CKKS homomorphic cryptosystem, and the exponents in Eq. (3) are computed homomorphically. The client decrypts the result and can obtain the label probabilities and hard predictions for each position. For the prediction, we use several numbers of regression coefficients, ranging from 8 to 64; as this number increases, both the obtained accuracy and the computational complexity increase (see Supplementary Table 5). We use a single parametrization of the cryptosystem (see the “EPFL-Lattigo team solution” section in the Supplementary document) for all the regression sizes, which keeps the cipher expansion asymptotically constant. The security of this solution is based on the hardness of the RLWE problem with Gaussian secrets.

#### SNU-CKKS

The SNU team applies one-hidden layer neural network for the genotype imputation. The model is obtained from Tensorflow module in plain (unencrypted) state, and the inference phase is progressed in encrypted stated for given test SNP data encrypted by the CKKS HE scheme. We encode each ternary SNP data into a 3-dimensional binary vector, i.e., 0 → (1, 0, 0), 1 → (0, 1, 0) and 2 → (0, 0, 1). For better performance in terms of both accuracy and speed, we utilize an inherent property that each target SNP is mostly related by its adjacent tag SNPs. We set the number of the adjacent tag SNPs as a pre-determined parameter *d*, and run experiments on various choices of the parameter (*d* = 8*k* for 1 ≤ *k* ≤ 9). As a result, we check that *d* = 40 shows the best accuracy in terms of micro-AUC. Since the running time of computing genotype score grows linear to *d*, the fastest result is obtained at *d* = 8. We refer the intermediate value *d* = 24 to the most balanced choice in terms of accuracy and speed.

The security of the utilized CKKS scheme relies on the hardness of solving the RLWE problem with ternary (signed binary) secret. For the security estimation, we applied the LWE estimator^55^, a sage module that computes the computational costs of state-of-art (R)LWE attack algorithms. The script for the security estimation is attached as a figure in the “SNU team solution” section in the Supplementary document.

### Non-Secure Methods

We describe the versions and the details of how the non-secure methods were run. The benchmarks were performed on a Linux workstation with 769 Gigabytes of main memory on an Intel Xeon Platinum 8168 CPU at 2.7 GHz with 96 cores. No other tools were run in the course of benchmarks.

#### Beagle

We obtained the jar formatted Java executable file for Beagle version 5.1 from the Beagle web site. The population panel (1,500 individuals) and the testing panel data are converted into VCF file format as required by Beagle. We ran Beagle using the chromosome 22 maps provided from the web site. The number of threads is specified as 16 threads at the command line (option ‘nthreads=16’). We set the ‘gp’ and ‘ap’ flags in the command line to explicitly ask Beagle to save genotype probabilities that are used for building the sensitivity versus PPV curves. Beagle supplies the per genotype probabilities for each imputed variant. These probabilities were used in plotting the curves.

#### IMPUTE2

IMPUTE2 is downloaded from the IMPUTE2 website. The haplotype, legend, genotype, and the population panels are converted into specific formats that are required by IMPUTE2. We could not find a command line option to run IMPUTE2 with multiple threads. To be fair, we divided the sequenced portion of the chromosome 22 (from 16,000,000 to 51,000,000 base pairs) into 16 equally spaced regions of length 2,333 megabases. Next, we ran 16 different IMPUTE2 instances in parallel, as described in the IMPUTE2 manual. The output from the 16 runs is pooled to evaluate the imputation accuracy of IMPUTE2. IMPUTE2 provides per genotype probabilities, which were used for plotting the precision-recall curves.

#### Minimac3

Minimac3 is downloaded from the University of Michigan web site. We next downloaded Eagle 2.4.1 phasing software. ‘Eagle+Minimac3’ is used in the Michigan Imputation Server’s pipeline that is served for the public use. The panels are converted into indexed VCF files as required by Eagle and Minimac3. We first used the Eagle protocol to phase the input genotypes. The phased genotypes are supplied to the Minimac3, and final imputations are performed. Both eagle and Minimac3 were run with 16 threads using the command line options (‘-numThreads=16’ and ‘-cpus 16’ options for Eagle and Minimac3, respectively). Minimac3 reports an estimated dosage of the alternate allele, which we converted to a score as in the above equation for UTMSR’s scoring.

## Acknowledgements

Authors thank the National Human Genome Research Institute (NHGRI) of National Institutes of Health for providing funding and support for iDASH Genomic Privacy challenges (R13HG009072).

We also thank Luyao Chen for providing technical support to set up the computational environment for unified evaluation.

## Author Contributions

MK, AH, and XJ designed the imputation scenario, implemented the evaluation metrics, conducted the benchmarking experiments with the baseline methods, and drafted the manuscript. MK, AH, XJ, YSG, and KL designed the baseline methods for UTMSR imputation pipelines. IC, SC, MG, and NG trained and implemented the Chimera-TFHE pipeline and contributed the results to the manuscript. DK, WC, SH, YSN, and JHC trained and implemented the SNU-CKKS pipeline. YM, JTP, DF, JPB, and JPH trained and implemented the EPFL-CKKS pipeline. HS, LOM, and XJ oversaw the iDASH19 challenge, conceived the study, and edited the manuscript. All authors have read and approved the final manuscript.

## Code availability

The source code implementation of the training and secure evaluation of the developed approaches will be made publicly available upon publication.

## Data availability

The array variants are available for download from Illumina’s web site. The 1000 Genomes Project datasets are downloaded from the NCBI ftp portal.

